# Image restoration of degraded time-lapse microscopy data mediated by infrared-imaging

**DOI:** 10.1101/2022.11.10.515910

**Authors:** Nicola Gritti, Rory M. Power, Alyssa Graves, Jan Huisken

## Abstract

Time-lapse fluorescence microscopy is key to unraveling the processes underpinning biological development and function. However, living systems, by their nature, permit only a limited toolbox for interrogation. Consequently, following time-lapses, expended samples contain untapped information that is typically discarded. Herein we employ convolutional neural networks (CNNs) to augment the live imaging data using this complementary information. In particular, live, deep tissue imaging is limited by the spectral range of live-cell compatible probes/fluorescent proteins. We demonstrate that CNNs may be used to restore deep-tissue contrast in GFP-based time-lapse imaging using paired final-state datasets acquired using infrared dyes and improve information content accordingly. Surprisingly, the networks are remarkably robust over a wide range of developmental times. We employ said network to GFP time-lapse images captured during zebrafish and drosophila embryo/larval development and demonstrate live, deep tissue image contrast.

## INTRODUCTION

Time-lapse imaging provides a uniquely dynamic view of biological processes in living systems ^1–6^. Powerful insights into development and function have followed through a union of modern microscopes and genetically encoded fluorescent proteins (FPs)^7,8^. Nevertheless, such an approach has its limits in terms of the type of information that can be extracted. For example, the ability to resolve deeply situated tissues in living animals or 3D cultures is circumscribed by the poor penetration of visible light therein, a challenge exacerbated by the need to maintain physiological conditions. However, in pursuit of a richer biological understanding, we should maximally leverage each specimen to extract complementary information that is so often left on the table as living samples are typically discarded following a time-lapse. For example, expended samples could exploit the less constrained toolbox available to fixed tissue imaging (multiplexed staining^9^/clearing^10^/expansion^11^ or even physical sectioning ^12^ or harsh imaging modalities that are less compatible with live imaging but may provide additional information of the specimens final state, such as those that use high illumination intensities ^13^, long recording times ^14^, harmful radiation ^15^, or restrictive mounting ^16^. Captured *in situ*, (i.e., in a single instrument), this approach can yield multimodal datasets that are useful unto themselves ^17^. An intriguing question is whether this additional information extracted from the final state, which is inaccessible to live imaging, can be leveraged to directly enhance the dynamic time-lapse data: ideally, merging the high spatial resolution data obtained at the final time point with the temporal information gained during the time lapse. Such an approach has been out of reach in the absence of a translation layer between the time-lapse and final state data. However, supervised machine learning approaches and in particular, deep neural networks are capable of learning complex, highly non-linear relationships between two associated datasets^18^ and have been applied to a range of bio-image restoration ^19–21^, segmentation ^22–24^, and classification tasks ^6,25^ Consequently, multi-modal microscopy and convolutional neural networks may be used to enhance *in vivo* time lapse microscopy images. The deep learning network could even be trained for cases featuring a single dataset characteristic of snapshots of the live and fixed states and subsequently applied to dynamic live imaging data.

As an illustrative use case, we consider the origin of image quality degradation deep in tissue. The poor penetration of visible light limits high-resolution fluorescent protein-based imaging to superficial regions in all but the smallest, most transparent embryos or isolated cells. Conversely, near infrared (NIR, 750 - 1700 nm) light maintains its directional propagation deeper into tissue ^26^, but suitable NIR fluorophores are incompatible with live imaging (a discussion of infrared imaging is given in Supplementary Note 1). Herein, we demonstrate *in vivo* time lapse microscopy with enhanced information-content on the basis of paired live (GFP) and final state (NIR) datasets. This technique, termed IR^2^ (infrared-mediated image restoration), is broadly applicable to a multitude of biological systems requiring only a green fluorescent protein (GFP) contrast for live imaging and a post fixation staining against GFP with NIR dyes. IR^2^ could thus provide a route to studies of biological dynamics in deeply located tissues and across later developmental stages than hitherto accessible.

## RESULTS

Restoration of contrast using IR^2^ requires a 1:1 correspondence between the endogenous GFP contrast (live and fixed) and the NIR staining (fixed). Firstly, the instrumentation must be simultaneously capable of fast and gentle imaging of the live-state and subsequent capture of the fixed state across a broad spectral region below and above 750 nm. We therefore developed a custom selective plane illumination microscope (IR-mSPIM, see Methods), capable of high-resolution imaging over this wide spectral range (see Supplementary Note 2, Chromatic performance and calibration of the IR-mSPIM). Although a light sheet microscope compatible with dyes emitting up to 1700 nm has been reported^27^, absorption from water increases > 100x compared to an optimum window at 800 - 950 nm^26^ contributing to heating and attenuation of the excitation/emission light. Furthermore, the deep cooled InGaAs cameras required to image beyond 1000 nm have unfavorable noise characteristics and a cost per pixel > 50x that of widespread silicon technologies. For these reasons and the desire to maintain performance in the visible, the IR-mSPIM achieves visible/NIR-I excitation at 488/640/808 nm and efficient collection at bands centered at 525/697/845 nm, alongside compatibility with moderate-high NA water dipping optics, suited to high-resolution live imaging.

Secondly, staining must be achieved with high-specificity and penetration into tissue without harsh treatments that would compromise/deform structures from the organismal level down to the resolution limit of the microscope. Notably, this latter point precludes the use of clearing protocols, which non-uniformly shrink or expand tissues^10^. Nevertheless, due to the diverse compositions of the different biological tissues and organisms, we are unaware of any staining protocol that preserves organismal structures and provides a homogeneous labeling for all tissue types. Rather, protocols need to be finely tuned specifically to each organism and transgenic line (for a description of the protocols used in this work, see the Methods section). This aspect should not be overlooked as, for instance, overfixation of the tissues may decrease the relative brightness of GFP and background autofluorescence^28^. Likewise, aggressive permeabilization is precluded by the need to maintain tissue structure/integrity down to the resolution level of the microscope, thus presenting limits to the passage of dye-tagged macromolecules. A discussion of the optimization of the protocols used in this work is provided in Supplementary Note 1.

Using IR-mSPIM, we first sought to demonstrate that infrared dye staining against GFP could be achieved with a high-degree of selectivity throughout mm-sized embryos/larvae. A fixed transgenic zebrafish larva expressing GFP in the vasculature (Tg(kdrl:GFP)) was imaged after immunostaining with AlexaFluor800 (AF800, ThermoFisher) (Figure 1 A and Supplementary Table 1). To perform an objective comparison of the visible and IR images, we extracted small volumes (patches of 128 × 128 × 32 pixels each) from the full images, and computed the pixel-wise Pearson correlation between the two (Figure 1 B). As we anticipate better depth penetration in the infrared, this metric is primarily influenced by the quality of staining as shown under conditions of poor selectivity (Supplementary Figure 1). As such, the R-value demonstrates that the staining proceeds with high selectivity and without increasing background. The vascular system is highly accessible to large percolating antibodies and so even staining of large samples (mm-sized) can be achieved quickly. We found denser, thicker tissues such as densely packed cell nuclei in the brain, to be more difficult to penetrate. As such we sought alternative staining strategies (see Supplementary note 3. Fixation, permeabilization and staining strategies). Compared to conventional antibodies, nanobodies are substantially smaller and thus potentially better suited in this regard ^29,30^. To explore whether nanobodies could be used to achieve homogenous staining of thick and dense tissues, we conjugated a GFP nanobody (Chromotek) with a NIR Cyanine-based fluorescent dye (CF800, Biotium) via maleimide chemistry and stained a zebrafish expressing a histone-GFP fusion Tg(h2b:GFP) (Figure 1 A’ and Supplementary Table 1). Selected patches and Pearson correlation demonstrated comparable selectivity to antibody staining (Figure 1 B’). While traditional immunostaining failed without a more destructive permeabilization (Supplementary Figure 2), even deeply located tissues showed uniform staining for this challenging case where the dye/nanobody must penetrate through multiple, mildly-permeabilized cell membranes and a nuclear envelope (Figure 1 C’). It is worth noting that both antibody/nanobody conjugation were associated with a strong redshifting of the excitation and emission spectra of the IR dyes compared to reference spectra with an expected further decrease in scattering.

**Figure 1:**
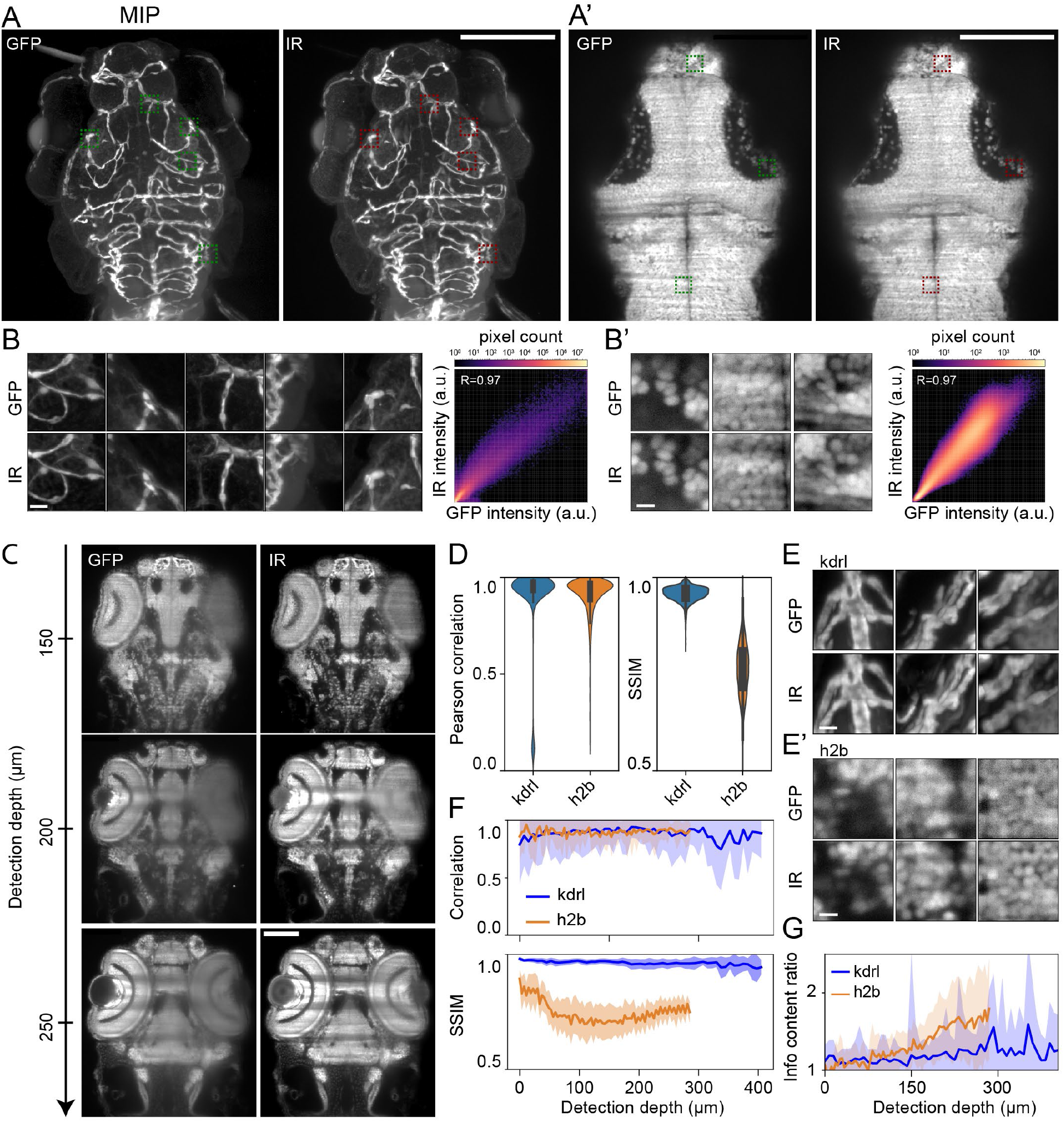
Highly-selective infrared staining and light sheet microscopy affords superior imaging at depth in tissue. **A)** Maximum intensity projected (MIP) z-stacks acquired for a fixed Tg(kdrl:GFP) (vascular marker) zebrafish larva (72 hpf) stained against GFP via conventional indirect immunostaining with AlexaFluor800 (AF800). Visible (GFP) left, infrared (AF800) right. Scale bar: 100um **A’)** A single superficial z-plane from a fixed Tg(h2b:GFP) (nuclear marker) zebrafish larva/embryo (96 hpf) stained against GFP via nanobody-conjugate CF800. Visible (GFP) left, infrared (CF800) right. Scale bar: 100um **B, B’)** Selected superficial patches shown by the dashed boxes in A, A’ respectively (visible (GFP) top, infrared (AF800/CF800) bottom) and pixelwise correlation plots for all 125 extracted patches. Scale bar: 5um **C)** Multiple deeper z-planes acquired for the same nuclear marker (h2b) zebrafish embryo/larva shown in B. Scale bar: 100um **D)** Pearson correlation and structural similarity index measure (SSIM) for the full z-stacks acquired for the vascular (kdrl) and nuclear (h2b) marker zebrafish from A, A’ respectively. **E)** Selected deeper patches for the vascular (kdrl) and nuclear (h2b) marker zebrafish from A, A’ respectively. Scale bar: 5um **F)** Pearson correlation and SSIM for all patches extracted at different z-planes from the full z-stacks of the vascular (kdrl) and nuclear (h2b) marker zebrafish. The z-depth provided is the maximum z-depth into tissue for each image in the stack. The uncertainty envelope is given by standard deviation. **G)** The information content gain (see Methods), between the visible (GFP) and IR channels (I_IR_/I_GFP_) for all patches extracted at different z-planes from the full z-stacks of the vascular (kdrl) and nuclear (h2b) marker zebrafish. The uncertainty envelope is given by standard deviation.

The improvement in image quality in the IR is clearly apparent at depth into tissue and provides a sound basis for image restoration. For both vascular and nuclear transgenic lines, the Pearson correlation clusters strongly towards 1, highlighting that staining is accomplished evenly throughout (Figure 1 D, note the tail towards lower values corresponds to patches dominated by noise i.e. dark regions of the image, Supplementary Figure 3). The structural similarity index measure (SSIM) for the vascular label is also close to 1 (Figure 1 D); the more deeply situated patches demonstrate that due to favorable feature sparsity and size one can follow individual vessels throughout even for GFP (Figure 1 A, E). However, for the nuclear marker, the majority of patches are clustered around a SSIM of ca. 0.75 (Figure 1 D). In this case, the deeply situated patches are notably different for the GFP and IR channels (Figure 1 E’). The depth-dependence of the SSIM demonstrates that the deviation between GFP/IR images occurs primarily at depth (Figure 1 F), while the near depth-invariance of the Pearson correlation again highlights its suitability to assess stain penetration. Since staining is achieved with a high degree of uniformity, the difference must arise from an improved image quality for the IR channel at depth. To obtain an absolute measure of image quality, we computed entropy-based information content as previously described ^31^(Methods), with the IR images at depth showing as much as 2.5× the information content of their GFP counterparts (Figure 1 G). We note that commercially-available nanobodies conjugated to the far-red dye AlexaFluor647 (excitation/emission peaks ca. 650/670 nm) performed comparably for staining and offered a more modest improvement to image quality at depth, with the advantage that more common hardware for imaging in the visible spectrum can be used.

Having demonstrated that the IR staining pipeline preserves structure while labeling uniformly throughout tissue depth and selectively for GFP, we considered whether a supervised deep learning approach ^32^could be used to restore a high contrast image from tissue-scattered GFP images. Image degradation resulting from scattering of a light sheet has been restored using complementary images from two opposed illumination directions ^33^. However, this method, proposed also by others^34^ remains limited in terms of depth penetration since the associated ground truth arises from comparably superficial regions, whereas tissues that are deeply situated with respect to both illumination directions remain inaccessible. In contrast, we use the superior IR images as a ground truth, thus attempting restoration of images degraded by scattering induced in both illumination and detection. To test whether this approach is generalizable to other model organisms, we used transgenic lines from both zebrafish and drosophila embryos where the cell nuclei are labeled with GFP (Tg(h2b:GFP) and Tg(His2AV-GFP), respectively), and stained them with GFP nanobody conjugated to the NIR dye CF800. We used these datasets in combination with a common convolutional neural network from the CARE package ^19^. First, we implemented an optimized patch extraction routine to minimize the amount of dark patches in the training dataset (Supplementary Figure 4) and to ensure that the IR and GFP patches are maximally aligned locally (Methods). Next, the restoration network was trained (see Supplementary Note/Methods) using the GFP and IR images of the nuclear marker as degraded and ground truth datasets, respectively. Upon application of the network to the degraded GFP data, restored images emerged with visibly enhanced contrast for both zebrafish and drosophila subjects (Figure 2 A, B and F, G), thus suggesting that the IR^2^ approach could be applied to two model organisms with distinct optical properties.

**Figure 2:**
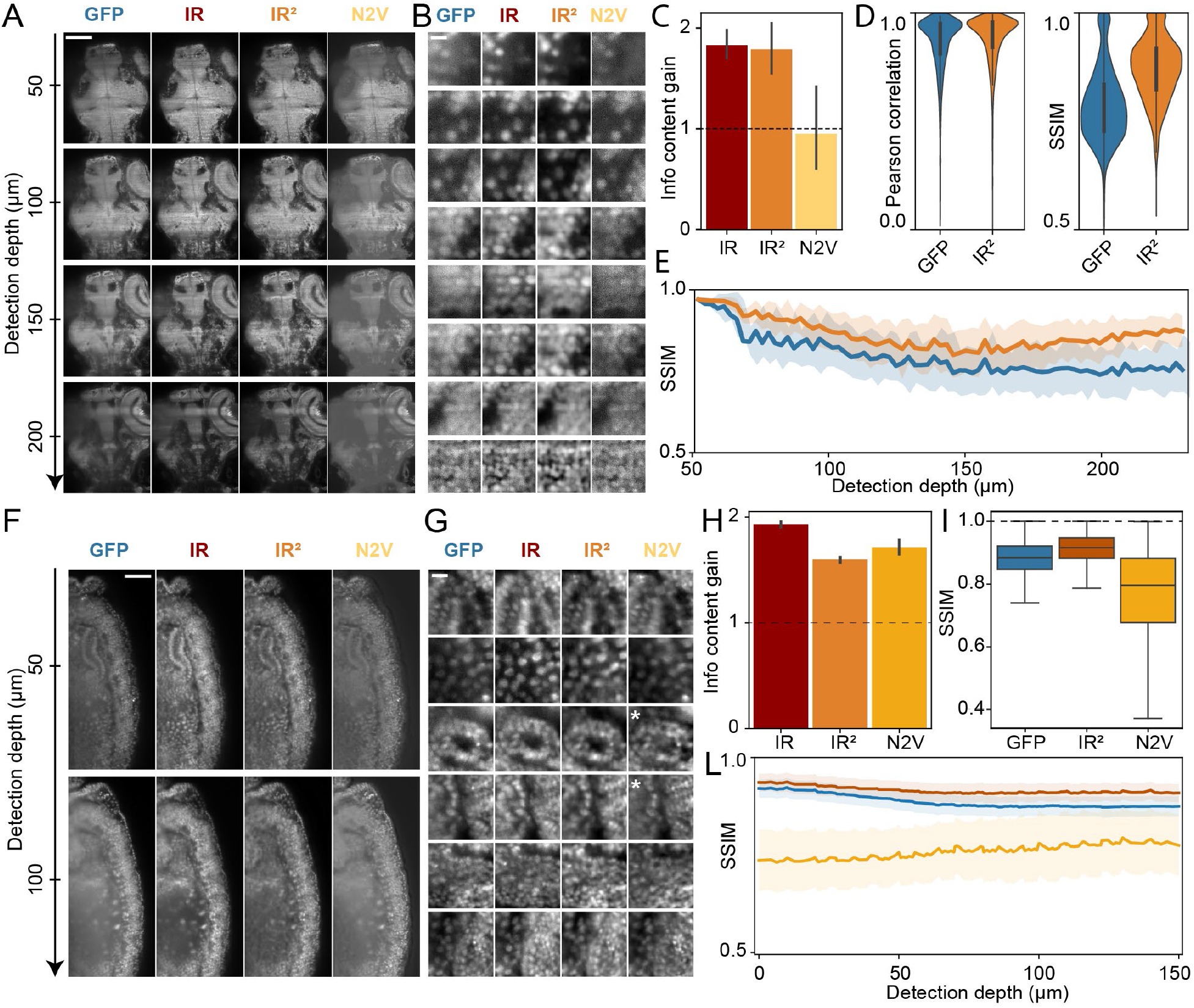
Infrared-mediated Image Restoration (IR^2^) improves image quality of degraded GFP images. **A)** Single GFP and IR images (left columns) extracted at increasing detection depth in a 96 hpf Tg(h2b:GFP) zebrafish larva restored with either IR^2^ or Noise2Void (N2V), (right columns). Scale bar: 100um **B)** Example patches for the same zebrafish dataset shown in A) arranged for increasing cell density. Scale bar: 5um **C)** Information content gain (relative to GFP) in patches extracted from the ground truth (IR, dark red), IR^2^-reconstructed (IRIR, orange) and N2V-reconstructed (N2V, yellow) images. **D)** Pearson correlation coefficient and global structural similarity index metrics (SSIM) obtained for patches extracted in the GFP and IR^2^-restored images when compared with the ground truth image (IR). **E)** Structural similarity index metrics (relative to IR image) as a function of detection depth for patches extracted throughout the sample. **F)** Single Z planes at increasing detection depth for a Tg(His2AV-GFP) fly larva (8 hpf) extracted from the input (GFP) and ground truth image (IR), as well as from restored images obtained from infrared-mediated and Noise2Void. Scale bar: 100um **G)** Example patches for the same fly dataset shown in F). White asterisks indicate patches where artifacts were introduced or features were not reconstructed by the Noise2Void network. Scale bar: 5um **H)** Information content gain (relative to GFP) in patches extracted from the ground truth (IR, dark red), IR^2^-reconstructed (IRIR, orange) and N2V-reconstructed (N2V, yellow) images. **I)** Structural similarity index metrics, relative to IR image, for GFP, infrared-mediated and Noise2Void reconstructions. **L)** Structural similarity index metrics as a function of detection depth.

We next set out to characterize the properties of the IR^2^ networks. To assess how IR^2^ images compare to a “blind” image denoising algorithm, we utilized Noise2Void (N2V), a deep learning approach capable of performing unsupervised image denoising by training on the GFP degraded images only^20^ (Figure 2 A, B and F, G). Notably, N2V images generated from zebrafish Tg(h2b:GFP) did not show an increased information content, suggesting that the degradation of the input GFP images was not dominated by noise (Figure 2 C). Instead, we observed that the IR^2^ images from the same input had significantly higher information content and approached that of the ground truth (IR), conversely suggesting that scattering at depth was the major source of degradation, and could efficiently be restored using IR^2^ networks (Figure 2 C). A similar behavior was observed for Drosophila GFP images, albeit the information content of the IR^2^ images did not reach the same levels of the IR ground truth (Figure 2 H). Notably, despite showing a similar information content gain, N2V reconstructions of Drosophila also displayed a lower structural similarity index (SSIM) with the ground truth (IR) (Figure 2 I), thus suggesting that while N2V networks may learn to efficiently denoise images, they were prone to introduce artifacts (Figure 2 G, white asterisks). The Pearson correlation of IR^2^ images, being primarily sensitive to the staining quality, remained similar across all patches, while the SSIM showed a remarkable improvement in the IR^2^ restored images, both for zebrafish and Drosophila (mean, SD = XX, XX respectively for GFP and restored images respectively), which is attributable to a contrast enhancement following restoration (Figure 2 D, I). The depth-dependent SSIM (Figure 2 E) showed an increasing deviation between the degraded (GFP) and ground truth (IR) images with depth. The SSIM decreased rapidly at first for the nuclear marker, highlighting that, even at modest depths, fine features in the image were substantially degraded. The restored data showed a much reduced degradation across all depths, highlighting the ability of the network to restore contrast lost due to scattering in tissue.

The success of deep learning-based restorative strategies relies on training data that is representative of the degraded data. When considering the restoration of dynamic time-lapse data, from static endpoint training data, one must consider the efficacy of the restoration over the full time series, over which substantial developmental processes may render morphological changes in the sample. To explore the effect of the developmental interval between the capture of live/degraded and fixed/ground-truth training data, GFP and IR images were captured of nuclear marker zebrafish (h2b:GFP) at 2, 3, and 4 days post fertilization (Figure 3 A). The GFP/IR from each age group were used to train restoration networks, each of which was subsequently applied to the task of restoring the degraded GFP images for all ages (Figure 3 B and Supplementary Figure 5). As expected, the network corresponding to the actual age produced the best restoration in all cases between the IR ground truth and GFP degraded data on the basis of normalized root mean square error (NMRSE) and SSIM (Figure 3 C). Nevertheless, the difference with the restoration based on dissimilar aged training data were small and all restoration networks resulted in an improved similarity with the IR images regardless of age. This suggests that the restoration networks can be successfully applied even over long developmental time windows, while a superior restoration of time-lapse data may be achieved by applying several trained networks over their respective developmental time window. We attribute this robustness to developmental time as a result of the origin of the signal to be restored, in this case, small-scale punctate features arising from cell nuclei (histones) largely dominating over large-scale morphological changes.

**Figure 3:**
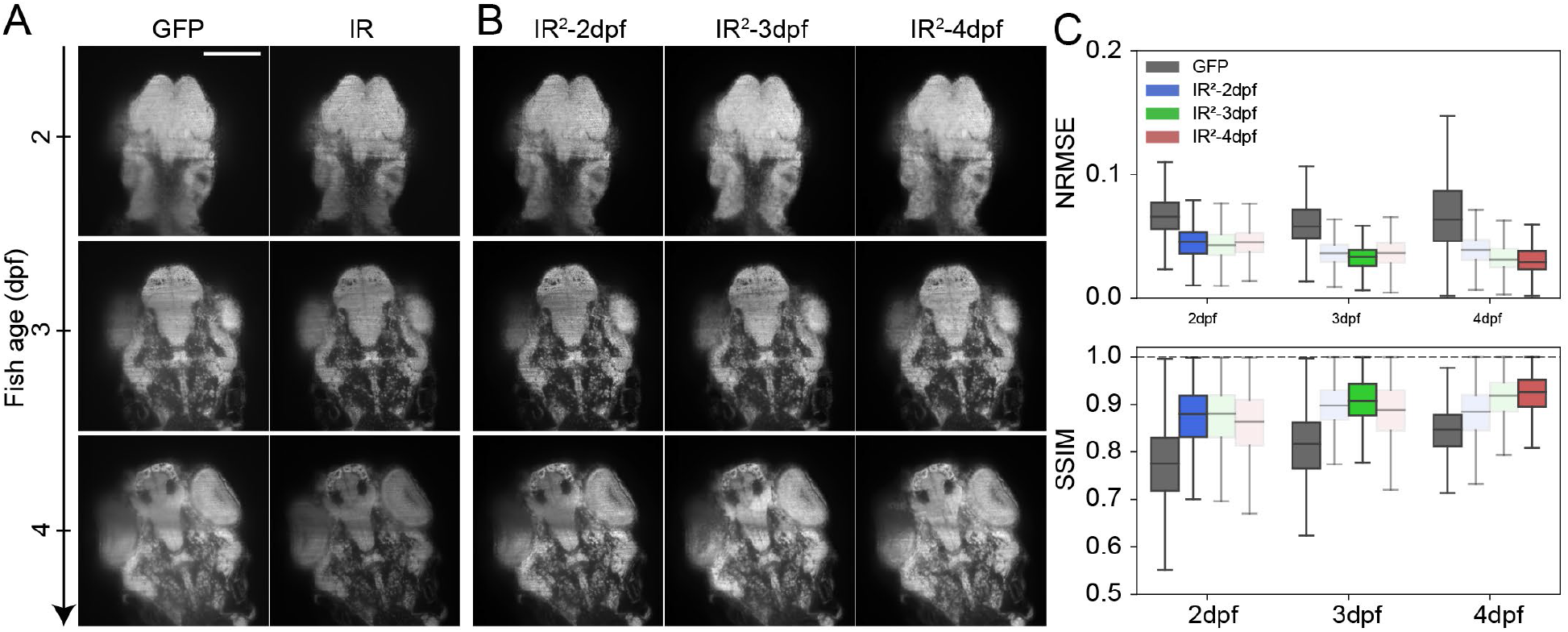
The quality of IR^2^ images is robust over large developmental intervals. **A)** Representative GFP and IR images of a single Z-plane in a full 3D stack of Tg(kdrl:GFP) zebrafish larvae at 2, 3 and 4 days post fertilization. Scale bar: 100um **B)** Image reconstructions obtained from IR^2^ models trained with images from 2, 3 and 4 dpf zebrafish larvae. **C)** Quantification of image quality as measured with the normalized root mean squared error (NRMSE) and structural similarity index metrics (SSIM) from the GFP image as well as the images obtained after IR^2^ reconstruction. Values were averaged over approximately 20,000 patches extracted from 5 different samples per age group.

Having demonstrated that infrared mediated image restoration performs well over wide developmental periods, we next set out to test whether this approach is applicable to continuous acquisitions of time-lapse live imaging data. To this end, we performed long term time lapse microscopy of developing zebrafish and drosophila embryos whose nuclei are labeled with GFP (Figure 4 A, E), and used IR^2^ networks trained on static images obtained via fixation and nanobody-staining of a sample. For both zebrafish and drosophila, we observed a qualitative increase in image contrast throughout the duration of the time lapse experiment (Figure 4 B, C, F and Supplementary Movie 1). To quantify the improvement, we computed the information content gain, defined as the information content of IR^2^ images relative to the information content of the corresponding GFP input for all z-slices and time points, (Supplementary Figure 6, Figure 4 G, H). We then computed the average information content gain in every plane of the z-stack and for every time point, thereby obtaining a kymograph representing the spatio-temporal evolution of the quality metric throughout the time lapse experiment (Figure 4 D, I). In both the zebrafish and the Drosophila cases, we observed an improvement in the information content gain with image detection depth. Conversely, the same quantity showed only a minor improvement with time, attributable to a general increase in the sample volume and so a relative improvement for the infrared imaging. This finding is consistent with the observations of Figure 3, that a single IR^2^ network maintains a similar restoration performance throughout a wide developmental range.

**Figure 4:**
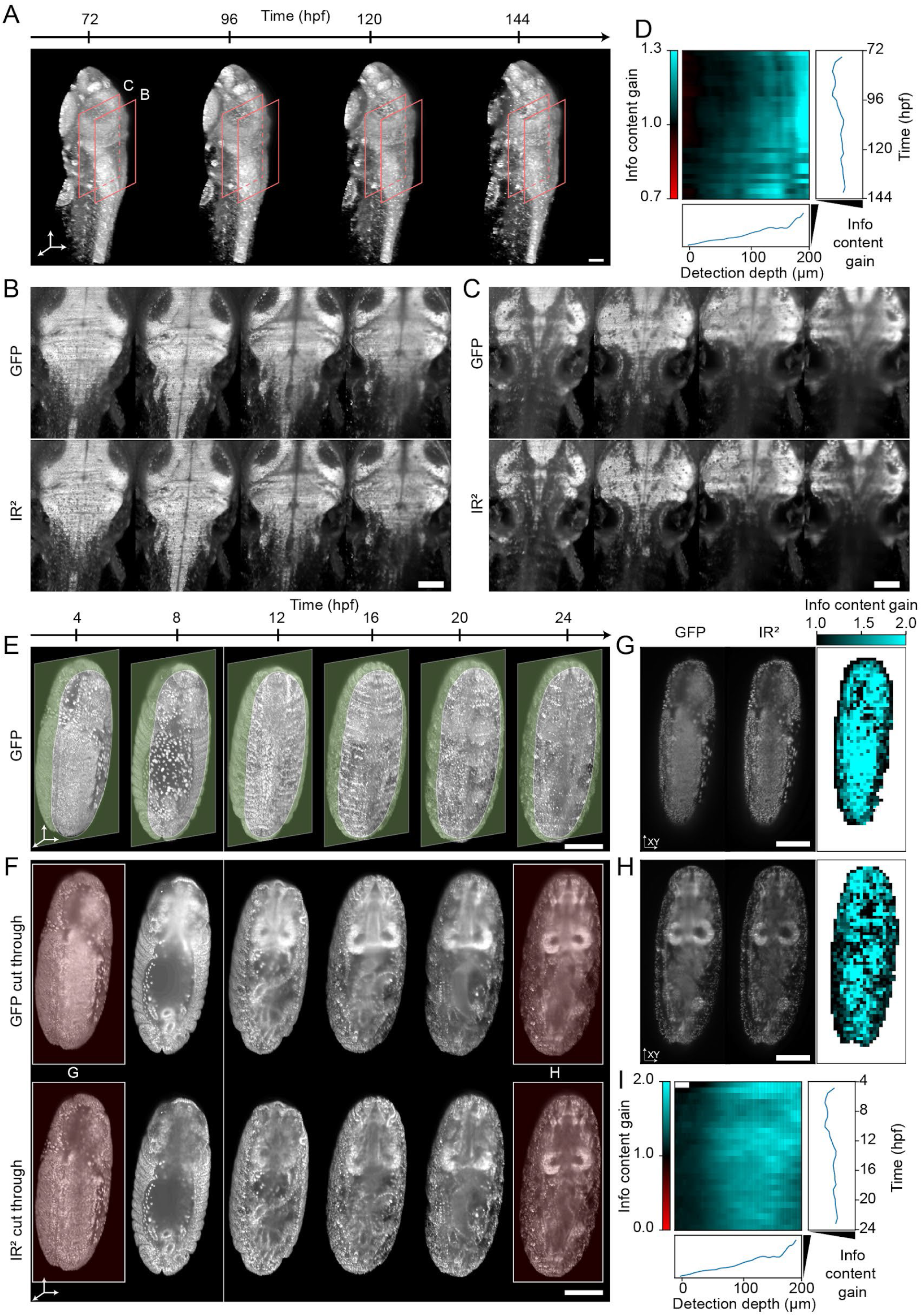
Infrared-mediated image restoration provides high-contrast deep-tissue time-lapse imaging of living biological systems. **A)** 3D reconstruction of live Tg(h2b:gfp) zebrafish larva images. Orange boxes represent the regions shown in panels B, C. Scale bar: 100um **B, C)** Individual Z-planes at detection depth=100um and 250um, respectively for the fish larva shown in panel A. Top row: endogenous GFP, bottom row: IR^2^ reconstructed images. Scale bar: 100um. **D)** Kymograph representing the information content gain relative to the GFP images, for all the images in the timelapse dataset and as a function of detection depth. Line plots to the right and the bottom represent the depth- and time-average information content gain. **E)** 3D reconstruction of live (Tg(His2AV-GFP)) fly larva images. Green opaque planes represent the sample sections shown in panel F. Scale bar: 100um. **F)** Individual Z-plane for the fly images shown in panel E. Top row: endogenous GFP, bottom row: IR^2^ reconstructed images. Red-highlighted timepoints are shown in panels G, H. Scale bar: 100um. **G, H)** Spatial mapping of the information content gain for the two individual Z-planes shown. Scale bar: 100um. **I)** Kymograph of the information content gain as a function of time and detection depth. Line plots represent the depth- and time-average information content gain.

Taken together, our results demonstrate that the information content of degraded time-lapse microscopy datasets containing only a visible contrast can be augmented by training a neural network on the basis of the relative benefits of deep-tissue infrared imaging and applying it to the task of image restoration to allow time-lapse imaging with high contrast even deep into tissue.

## DISCUSSION

We have demonstrated that supervised deep learning can be used to restore image quality in deep tissue images given suitable ground truth and degraded images. As a first illustration of this concept, we introduce IR^2^, which exploits infrared dye labeling of GFP as a route to paired degraded and ground truth datasets, alongside light sheet microscopy to provide fast and gentle live imaging. In contrast to existing restorative deep learning pipelines such as CARE and Noise2Void, which, while powerful in their own domains are not designed to enhance depth penetration in optical imaging, IR^2^ has been developed specifically to restore deep-tissue contrast to live imaging data from zebrafish and drosophila embryos/larvae. En route, we have demonstrated the utility and robustness of this approach to resolve features of embryonic/larval development across wide developmental time windows and which would otherwise be inaccessible owing to the limited penetration of visible light into tissue.

The methods reported can be generalized, requiring only widespread GFP lines and some optimization of staining protocols. Even in the absence of a specialist microscope capable of visible-IR imaging, improvements to image quality can likely be made for commercially available GFP-nanobody tagged dyes that are efficiently excitable in the visible range of the spectrum and an appropriate microscope (Supplementary Figure 2). The deep learning networks themselves require only modest computational resources. In this regard, hardware requirements, software, and pre-trained models/datasets are provided to aid uptake of these restorative abilities by biologists seeking to perform minimally invasive live deep tissue imaging (GitHub link).

A potent direction for the future would be to explicitly incorporate a depth dependent component to the restoration, using the detection depth as an additional channel. Furthermore, model training has been carried out using only single samples, rather than by combining ensembles. Limited training data is a general challenge to deep learning methods, however, light-sheet techniques are able to generate vast quantities of data rapidly. Since no annotation is required, several datasets could be combined to learn additional features for restoration at the cost of increased training time.

We expect further improvements to the performance of IR^2^ commensurate with developments in fixation protocols that better maintain tissue structure and GFP fluorescence. Similarly, more photostable and brighter IR dyes, redshifted towards the short wave infrared^35–37^ (alongside commensurate developments in low noise cameras) may allow even deeper tissue imaging. Furthermore, for widely distributed/shared technologies^38^, a library of images and restoration networks could be curated for given transgenic lines and shared with other users thus allowing restoration to be applied as an optional part of their post-processing pipeline.

The IR^2^ restoration network is not limited to the basis of GFP/IR images as degraded/ground truth pairs and could prove similarly powerful for ground truth images arising from the use of adaptive optics, multiphoton excitation or indeed chemical clearing if tissue distortions can be obviated. In pursuit of minimizing animal usage, the scheme outlined provides one route by which the information contained in a single subject may be additionally leveraged rather than lost when discarding samples after time-lapse imaging. We anticipate that IR^2^ can provide a powerful tool in the biologist’s arsenal for deep tissue live imaging.

## Acknowledgments

We would like to thank the entire Huisken Lab for fruitful discussions on this topic. N.G. was supported by the Human Frontier Science Program (HFSP) fellowship (LT000227/2018-L) R.M.P was supported by the Human Frontier Science Program (HFSP) fellowship (LT000321/2015-C; R.M.P.) and J.H. was supported by the Morgridge Institute for Research. We thank the lab of Jill Wildonger for assistance with drosophila husbandry/handling, and provision of transgenic animals. We thank the Biomolecular Screening and Protein Technologies Unit (CRG, Barcelona) for assistance in the nanobody conjugation.

## METHODS

### Zebrafish husbandry and transgenic lines

Zebrafish (Danio rerio) were handled according to established protocols approved by the University of Wisconsin-Madison Animal Care and Use Committee. Zebrafish adults and larvae were maintained on a 14h/10h light/dark cycle at 28 °C. Zebrafish embryos were raised in E3 media at 28 °C. Larvae raised beyond 5 days post fertilization (dpf) were fed a normal diet until commencement of imaging (powder mixture of 1/3 Spirulina, Encapsulon, and ground Tetramin Tropical flakes). Transgenic lines Tg(kdrl:GFP) and Tg(H2b:GFP) were outcrossed to a casper background to reduce pigmentation where possible. Phenylthiourea (PTU) was used for depigmentation otherwise. Individual positive embryos were chosen randomly from a clutch of 100 - 300 embryos (at a density of ~0.5 fish/ml) or from pooled clutches where necessary.

### Fixation and Immunostaining of Zebrafish

Standard protocol: The standard protocol was employed for fixation/staining of transgenic lines that did not show appreciable limitations to penetration of antibodies. Embryos/larvae were fixed in 1.5% paraformaldehyde (PFA) in phosphate-buffered saline (PBS) + 0.5% Triton (PBST) for 2h at 4 °C and then washed overnight in aldehyde block (0.3 M glycine in PBST) at 4 °C. The fixed fish were briefly washed in aldehyde block before being permeabilized in PBST for 4h at room temperature (RT). Subsequently, fish were washed for 1, 2, 5, 30min RT in PBST and blocked for 2h RT in 0.05 % Tween, 0.3 % Triton, 5 % normal goat serum (NGS), 5 % bovine serum albumin (BSA), 20 mM MgCL_2_ and PBS. After a brief wash in PBST, fish were incubated consecutively overnight at 4°C and 2h at RT in primary and secondary antibodies respectively (diluted 1:500) in PBST + 5 % goat serum. Finally, embryos/larvae were washed in PBS until ready for imaging. In the case of nanobody staining, after the blocking step, fish were incubated for 2h at 4 °C (diluted 1:500 or 1:100) and then washed in PBS until ready for imaging.

Trypsin protocol: The trypsin protocol was carried out for fixation/staining of transgenic lines for which antibodies failed to penetrate tissue when using the standard protocol, the rationale being that a more aggressive permeabilization with trypsin could aid penetration. Embryos/larvae were fixed and washed overnight following the standard protocol. Next, the fish were permeabilized in 0.25% trypsin in PBS for 5 min on ice, washed briefly in PBST and continued from the blocking step from the standard protocol to completion. (protocol modified from ^39^) The protocol was not effective in enhancing penetration of the antibodies (see Supplementary Figure 2).

### Zebrafish mounting

Live embryos and larvae were first anesthetized in E3 media (without methylene blue) containing 0.16 mg/ml Tricaine (Sigma) and embedded for imaging in 2 % low-melting-point agarose/E3 (Sigma) with an internal diameter (ID) of 0.8mm and outer diameter (OD) of 1.2mm (Proliquid). Imaging was carried out at room temperature and the chamber was filled with RO water for fixed samples and-tricaine/E3 media for live samples.

### Drosophila husbandry and transgenic lines

Fly stocks were maintained by the lab of Jill Wildonger at the University of Wisconsin-Madison according to established protocols approved by the University of Wisconsin-Madison Animal Care and Use Committee. Flies were kept on a 12h/12h light/dark cycle and transferred to fresh vials with food every two days. The Tg(His2AV-GFP) transgenic line was used.

### Fixation and Immunostaining of Drosophila

Embryos were collected at the desired developmental stage, rinsed in RO water, and placed for 90 seconds in a petri dish with 100% bleach to weaken the outer shell. A paint brush was used to roll the embryos on the petri dish surface and remove the shell before rinsing with RO water. Embryos underwent a first fixation of 1:1 of 9% PFA in PBS: heptane for 30min RT. The inner vitelline membrane of embryos was removed by filling the embryo-containing vial with 55 % heptane and 45 % methanol and striking the vial against a table surface for 2 min, settling for 2 min, and repeating for a total of 3 times. Supernatant and floating non-cracked embryos were removed and an aldehyde block (0.3M glycine in PBST) was added for an overnight incubation at 4 °C. Next, embryos underwent a second fixation with PBST for 4 h at RT, washed with methanol and then ethanol, and blocked with 0.3 % Triton, 3 % BSA, 10 mM glycine, 1 % goat serum, 1 % donkey serum, and 2 % DMSO in PBS at 4 °C overnight. The embryos were incubated in primary antibody (1:500) overnight at 4 °C in PBS + 5 % goat serum, washed twice, and then stored in wash buffer overnight at 4 °C (consisting of 0.1% Triton, 3% BSA, 10 mM glycine in PBS and adjusted with NaOH-HCl to pH 7.2. After a brief wash of PBST, the embryos were subsequently incubated for 2 h at RT in secondary antibody solution (1:500) in PBS + 5 % glycine. When using nanobodies, after the blocking step the embryos were incubated for 2 h at 4 °C (1:500 or 1:100). After antibody/nanobody incubation, embryos were washed in PBS until ready for imaging. (Protocol modified from^40^).

### Drosophila mounting

Live and fixed embryos were embedded for imaging in 2 % low-melting-point agarose/PBS in FEP tubes with an ID of 0.8mm and an OD of 1.2mm (ProLiquid). Imaging was carried out at RT and the chamber was filled with RO water for fixed samples and PBS for live samples. A number of embryos were mounted in each tube to identify suitably oriented candidates for imaging (with their body axis approximately aligned along the tube axis).

### IR-mSPIM

Visible and infrared excitation was provided by a Toptica MLE laser engine (SM-fiber-coupled: 405 nm, 488 nm, 561 nm, 640 nm all 50 mW) and Omicron, LightHub-4 laser combiner (free-space: LuxX: 685 nm, 50 mW, 785 nm, 200 mW, 808 nm, 140 mW). The collimated laser outputs were expanded in one dimension using pairs of cylindrical lenses. The visible and infrared lasers were combined via a shortpass dichroic mirror. The light sheets were produced by cylindrical lenses, using a galvo mirror-based (Scanlab Dynaxis 3S) mSPIM scheme to pivot the individual light sheets for efficient stripe suppression^41^. Ultra-broadband achromatic doublets (400 - 1000 nm) were used where possible to relay and deliver the light sheets into a sample chamber with coverslip windows via two opposed water-corrected air immersion illumination objectives (Zeiss, LSFM 10x/0.2)

The emission path was optimized for visible-NIR transmission. A multiphoton objective (Olympus XLPLNS10XSSVMP 10x/0.6, 8 mm WD) provided good transmission and moderate NA with a large field of view. Nevertheless, axial chromatic aberration at < 650 nm required correction via automatic refocusing of the lens and immersion chamber using a motorized stage Physik Instrumente, M-111.1DG) (typical change in focal plane +/- 10 um). The light sheet remains at a single z-plane throughout refocusing and as such the location of the imaged plane does not change as the objective and chamber are moved. The chromatic calibration procedure and performance are discussed in Supplementary Note 2. A tube lens (400–1300 nm, Thorlabs TTL200MP, 200 mm focal length) was used to produce an image of the sample at 11.1x magnification on an sCMOS camera (Andor Zyla 4.2) with non-negligible sensitivity up to ca. 950 nm. The magnification of the system could be increased to 22.2x by exchanging the tube lens with an ultra-broadband achromatic lens (Thorlabs, AC508-400-AB-ML). Fluorescence was spectrally filtered from the excitation using bandpass filters (Chroma ET525/50m, ET697/60m, ET845/55m for GFP, AlexaFluor 647, AlexaFluor 800/CF800 respectively) mounted in a motorized in a filter wheel (Ludl 96A351, MAC6000 controller). Samples were mounted in FEP tubes via a custom sample holder. 3 translation stages and one rotation stage (Physik Instrumente M111.1DG, U-651 with C-884, C-867 controllers) were used to orient the sample and acquire z-stacks. Hardware control and synchronization were provided by custom LabVIEW software and a USB-6343 Multifunction DAQ device (National Instruments).

### Nanobody conjugation

The anti-GFP VHH/ Nanobody (Chomotek) was site directed conjugated with the CF680R maleimide (Biotium, Hayward, CA). The nanobody at concentration of 100uM was incubated for 2 h at room temperature with an equimolar amount of dye. Labeled protein was separated from unlabeled protein by size exclusion chromatography.

### Deep Learning

Upon acquisition of the GFP images and their IR counterparts, we obtained training samples by generating patches of dimension 128×128×32 pixels throughout the Z stack. Patches were extracting using either homogeneous distribution, that is, using a probability of extraction per pixel equal to:

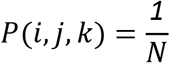

Where N is the total number of pixels in the Z stack, or using a selective probability:

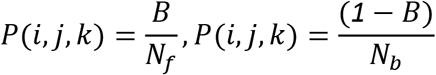

In this case, a threshold was computed using the Otsu thresholding and pixels were classified as foreground (pixel value higher than threshold) or background (pixel value lower than threshold). Nf and Nb represent the total number of foreground and background, respectively. B is a tunable parameter used to adjust the fraction of patches extracted in foreground regions, where a value of B=Nw/Nd corresponds to the homogeneous probability distribution. Throughout the experiments, we used B=0.9, thus including only 10% of background patches in the extracted training dataset. Sample coverage was iteratively monitored by comparing the number of foreground and background pixels extracted in the training set, and patch extraction was interrupted when sample coverage reached a value of 95% (Supplementary Figure 4).

To avoid a misalignment between input and ground truth datasets due to residual chromatic aberrations, we subsequently performed a correlation based registration using local translations of the patches and found the (dx,dy,dz) translation that maximized the functional:

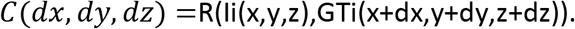

Where R represents the image cross correlation function, and Ii, GTi represent the input and ground truth patch, respectively.

With the training dataset thus obtained, we subsequently trained a deep learning network using the CARE framework^19^. In particular, throughout all experiments, we used a UNet algorithm with one-channel input and one-channel output, two hidden layers and softmax output layer (Supplementary Figure 4). The weights of the network were iteratively updated at every epoch using the mean squared error computed between the output of the network and the IR ground truth as a loss function. The input GFP image is thereby transformed at every subsequent layer into a new image with decreased spatial dimensions and increased channel dimension. Patches were divided into training and validation datasets with a ratio of 9:1, and the networks were trained over 100 epochs using a batch size of 8. Depending on the number of patches, training lasted approximately 12-24 hours using a GPU Quadro P5000 (16GB Memory) on a CentOS system (512GB RAM). Prediction of new images was performed on the same computational setup. All subsequent CPU-based image-based analysis, such as image information content, structure similarity index and root mean square error, were parallelized to use the 80 cores available.

### Entropy-based information content

Image information content is measured by computing the Shannon entropy of the discrete cosine transform of the image patch:

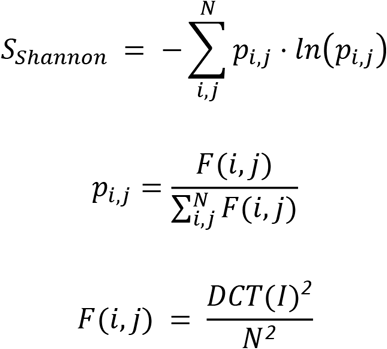

Where N represents the size of the patch and DCT is the discrete cosine transform of the image patch I.

The information content gain of an image 1 relative to an image 2, is defined as the ratio between the information content values of the two images.

## Data availability

A sample of the data and the code used are available on the Zenodo repository (doi: 10.5281/zenodo.7075414). Full datasets are available upon request.

