## Supplementary material for "Image restoration of degraded time-lapse microscopy data mediated by infrared-imaging"

### Supplementary Note 1: Biological infrared imaging.

GFP is the most widespread fluorescent contrast in live imaging. Nevertheless, the directionality of visible light is rapidly lost in tissue due to scattering, degrading the contrast of common microscopes beyond a few tens of µm depth. For example, the workhorse of 3D biological imaging, the laser-scanning confocal microscope functions well within this range only as scattered photons are rejected by the pinhole and ballistic photons are too few to generate meaningful contrast. Since scattering scales as an inverse power law with wavelength, longer infrared wavelengths are more able to penetrate deeper into tissue while maintaining their directional propagation. For example, optical coherence tomography is a reflection based imaging technique that exploits this superior penetration but does not yield a chemically specific fluorescent contrast.

Owing to its non-linear operation at infrared wavelengths and crosstalk-free point detection, multiphoton microscopy allows comparatively deep tissue imaging at several hundreds of µm depth while using visibly emitting fluorophores. While this is sufficient depth penetration for *in toto* imaging of small animal models such as embryonic/larval zebrafish and drosophila, these delicate samples are damaged by the intensely pulsed light and develop too quickly for their dynamic processes to be captured via serially point-scanned schemes.

As such, light sheet fluorescence microscopy (LSFM) has been the tool of choice owing to a powerful blend of parallelization and intrinsic optical sectioning, allowing fast, high-resolution biological imaging with low photodamage. However, the challenge of imaging deep into tissue remains. Furthermore, since image formation is dependent on both excitation and emission light, analogous multiphoton methods, which address the former only, do not offer the same tissue penetration as their point-scanned analogues[2]. To image deeper with LSFM, both the excitation and emission must be shifted to the near infrared (NIR-I, 700 -1000 nm).

Neither FPs nor live-cell dyes extend fully into the NIR-I and each is beset by unique challenges. Infrared FPs (IRFPs, with far-red excitation) suffer from being dim, weakly photostable, often dimeric and require biliverdin as a chromophoric co-factor[3]. Bioorthogonal schemes for targeting of live cell dyes such as Halo/SNAP- tagging systems require the expression of the self-labeling protein, and retaining cell-permeability places constraints on the dye redshift currently achievable[4]. Furthermore, neither tagging systems nor IRFPs are widely available in animal models and using more widespread biochemical tools is preferable. Under fixed tissue conditions, immunofluorescence provides a simple route available for targeted labeling of proteins, while non-permeant NIR-I dyes are widely available.

#### Supplementary note 2. Chromatic performance and calibration of the IR-mSPIM.

The IR-mSPIM corrects axial chromatic aberrations (i.e. a wavelength dependent working distance) by motorizing the detection objective (optimized for multiphoton imaging in the NIR) and immersion chamber. The light sheets are launched by air objectives, through chamber windows and so the light sheet position and hence imaged section remains static when refocusing in this manner. The correct refocus position was determined by imaging fluorescent bead phantoms dispersed in agarose. A mixture of beads was used to allow excitation at all available laser wavelengths (405, 488, 561, 640, 685, 785, 808 nm). The phantom comprised TetraSpeck (excitable at 405, 488, 561 and 640 nm) and Degradex PLGA NIR (excitable at 640, 685, 785 and 808 nm) fluorescent beads. The correction thus determined corresponds to the pass-band of each emission filter rather than the laser line used for excitation. As such, the correction is valid for other fluorophores (e.g. GFP, AlexaFluor dyes) when using the same emission filters. The optimum working distance was determined from the maximum variance achieved for a defocus series for each laser/emission filter combination. The cross-excitability of the two fluorescent bead species used provided the basis for aligning the far-red/NIR lasers to the visible lasers (which are inherently co-aligned out of the single-mode fiber): The far-red/NIR lasers (685, 785 and 808 nm) are aligned by visualizing the same subset of beads as when using the 640 nm line, using the same emission filter (845/55 bp) in each case, hence decoupling the laser alignment from the chromatic aberration of the objective lens. Any residual misalignment of the individual light sheets is accounted for via a registration step (see “Deep Learning” in

Methods). Lateral chromatic aberrations (changes in magnification with wavelength) are negligible and so the refocus scheme is sufficient to correct for the dominant chromatic aberrations.

The fluorescent beads are sub-diffraction sized providing a route to explore the chromatic performance of the IR-SPIM via the PSF. The scaling of lateral spatial resolution with wavelength would suggest that the performance is highly dependent on the specific emission band of the fluorophore and hence that the IR dyes would provide about half the resolving capability of GFP. However, SPIM often employs undersampling in the imaging path (with respect to NA), to allow for a larger field of view, while maintaining the light collection and axial resolution benefits of high NA. The data presented are captured at ca. 11.1x and 22.2x. The magnification can be switched between these two values by exchanging the tube lens (effl = 200 or 400 mm for 11.1, 22.2x respectively when paired with an Olympus 10x objective, effl = 18 mm). In the former case, given a pixel size of 6.5 um, the Nyquist-Shannon criterion provides a sampling determined resolution limit of ca. 1.2 um. Using the Rayleigh criterion for resolution:

res=0.61.lambda_0/NA

And taking lambda_0 in each case as the pass-band centre (525, 845 nm respectively for GFP, AF800/CF800), the expected resolution assuming diffraction-limited performance at NA = 0.6 is ca. 534, 859 nm respectively, both of which are below the Nyquist-Shannon criterion, and so it is sampling rather than wavelength that determines the spatial resolution. However, in the latter case, the Nyquist-Shannon criterion provides a sampling determined resolution limit of ca. 0.6 um, which could allow diffraction limited resolution for wavelengths > 590 nm and a substantial difference in resolution for GFP, AF800/CF800 respectively. The assumption made is that the imaging system performs in a diffraction-limited manner over the full wavelength range. The illumination system may be safely disregarded for consideration of lateral resolution however, the imaging system formed by the imaging objective lens (Olympus XLPLNS10XSSVMP 10x/0.6, 8 mm WD) and ultra-broadband tube lens (Thorlabs, TTL200MP/AC508-400-AB-ML) may not provide diffraction limited resolution, particularly the objective, which is optimized for two-photon imaging (most commonly ca. 920 nm). Given that axial chromatic aberrations are substantial for wavelengths < 700 nm, it is reasonable to also expect a decrease in the effective NA for shorter visible wavelengths and a convergence in the resolution for GFP and CF800/AF800. To explore whether this was apparent, bead stacks were analysed using PSFj[5]. The lower bound for the lateral PSF FWHM for excitation/emission centers of 488/525 nm, 640/697 nm, 808/845 nm was found to be 741 ± 14, 797 ± 15 and 974 ± 18 nm, which are equivalent to maximum NAs of 0.44, 0.54, 0.54 respectively (from the Rayleigh criterion), demonstrating that the optical performance is best in the far-red - NIR as expected and approaches the theoretical value of 0.6. Although the lateral PSF for the GFP equivalent is slightly narrower, we note that this will be apparent only in extremely superficial regions. In any case, the difference is minor and cell nuclei (ca. 5 - 10 um), which are the smallest structural details that we sought to resolve were easily resolvable for GFP, AF647 and CF800. The axial PSF FWHM is also similar for the three imaging bands: 5.46 ± 0.22, 5.36 ± 0.25 and 6.38 ± 0.41 um for 488/525 nm, 640/697 nm, 808/845 nm. In future, $IR^2$ could be combined with deconvolution strategies to improve the axial resolving power of multi-view light sheet microscopy as necessary[6].

### Supplementary note 3. Fixation, permeabilization and staining strategies.

The deep learning-based restoration requires as close to 1:1 correspondence between i) the live and fixed state of the animal and ii) the distributions of GFP and the infrared dye therein post-staining. Several key challenges are apparent. Firstly, the fixation should maintain GFP fluorescence and not enhance autofluorescence. Secondly, the permeabilization step should allow antibody penetration but not distort or degrade the sample morphology. Thirdly, the staining step should allow permeation of the antibodies throughout the tissue to evenly stain without non-specific binding.

To ensure these requirements were met, various approaches for fixation, permeabilization and staining of the tissue were tested and optimized for zebrafish samples. More generally, PFA fixation was found to be suitable for maintaining GFP fluorescence in all cases presented, however, the quenching of aldehydes via glycine washing reduced fixation-induced autofluorescence substantially. Various permeabilization steps were explored including organic solvents (methanol/acetone) and nonionic surfactants (Tween/Triton/DMSO) and proteinases (trypsin/proteinase K). Specifically, we tested reported protocols for zebrafish staining with slight modifications[1,7,8]. In some cases, antibody staining was lengthened to 7 days primary, 7 days secondary in an unsuccessful attempt to improve penetration. We found the standard protocol discussed in the methods section was sufficient for penetration of the vasculature label (Tg(kdrl:GFP) and best preserved the structure and gave good labeling-fidelity for a number of dyes. Nevertheless, the results from some dyes/affinity-tags were better than others. A large number of IR dye candidates were tested and protocols utilizing primary antibodies only, primary and secondary antibodies, as well as nanobodies were assessed. Note that there are many important characteristics of a dye and affinity-tag such as brightness/photostability, excitation/emission maxima, solubility, specificity (IR dyes are typically large lipophilic molecules bearing several charged functional groups that offer water-solubility at a potential cost to specificity), staining time required and completeness of staining. It was not possible to complete a full combinatorial assessment of all dyes and affinity tags independently, rather we were limited by the commercially available options and cost constraints. The dyes, antibodies and nanobodies tested that result in inferior (dimmer/less specific) staining than the case presented for the transgenic vasculature line Tg(kdrl:GFP) in Figure 1 is given in Supplementary Tables 1 and 2. We note, other lines, permeabilization and staining strategies or model organisms may provide better results with these products. None of the protocols attempted provided anything more than superficial penetration in the nuclear label (Tg(h2b:GFP) when used with antibody labeling. The custom-labeled CF800 nanobody (see Methods section), however, provided excellent staining in zebrafish and drosophila alike.

**Supplementary Table 1.** Combination of primary, secondary antibodies and dyes used throughout.

| Primary | Manufacturer (Cat. number) | Secondary | Manufacturer (Cat. number) | Notes | Fig. |
|---|---|---|---|---|---|
| aGFP | ThermoFisher (A-11122) | AF800 | ThermoFisher (A-32808) | Primary-secondary incubation | Fig. 1 (kdrl:GFP), Sup. Fig. 2 |
| nGFP | | AF647 | Chromotek (GB2AF647) | Commercially conjugated | Sup. Fig. 2 |
| nGFP | Chromotek (GT-250) | CF800 | Biotium (#92128) | Custom conjugated | Fig. 1 (h2b:GFP), Fig.2, Fig.3, Fig.4 |
| nGFP | Chromotek (GT-250) | AF700 | ThermoFisher (A-21038) | Custom conjugated | Sup. Fig. 1 |

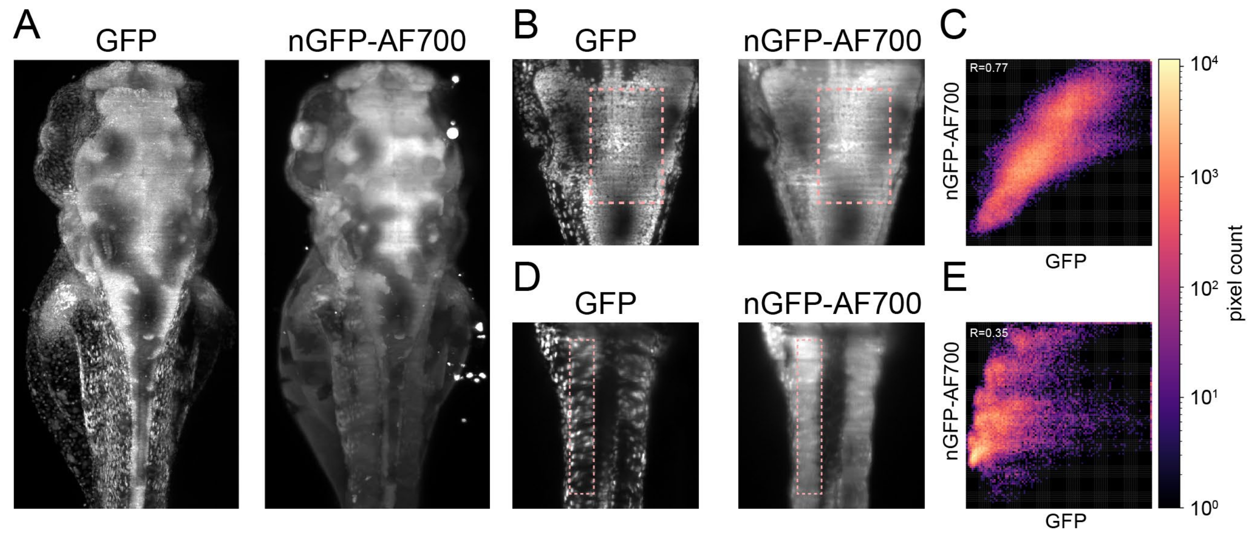


**Supplementary Figure 1.** A) Maximum intensity projection of a Tg(kdrl:GFP) zebrafish larva at 3 days post fertilization, stained using a nanobody conjugated to a dye (Alexa Fluor 700) in the NIR regime. B, D) Maximum intensity projection of a substack of the same fish in panel A) in the brain (B) and tail (D) region. C, E) Pixel-wise Pearson correlation coefficient for the regions highlighted in panels B and D.

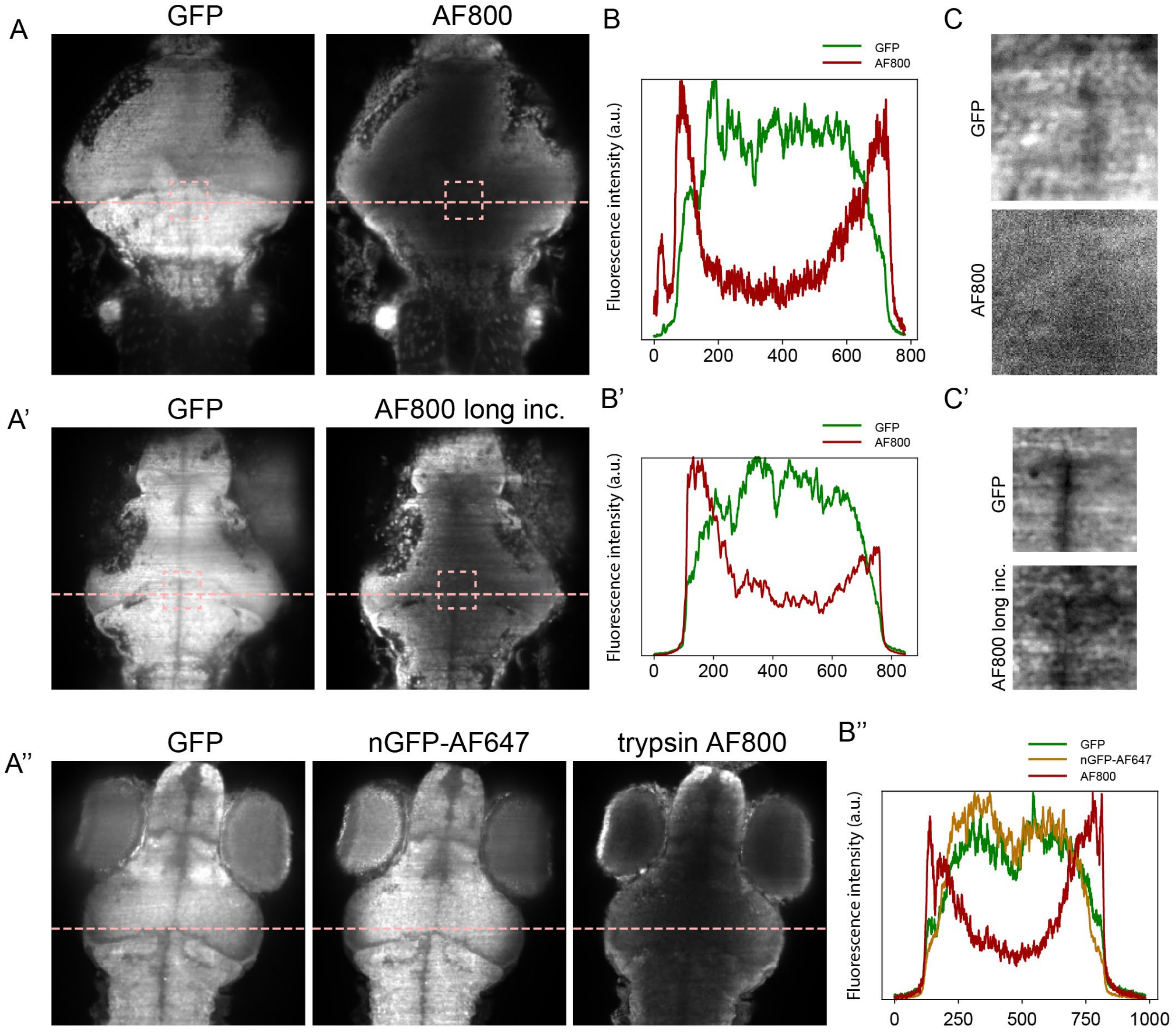


**Supplementary Figure 2.** A,A',A'') Single focal plane images from Tg(h2b::GFP) zebrafish stained with conventional primary-secondary antibody protocol (A), antibody protocol using 7 days incubation period (A') and simultaneous nanobodies and trypsin-based antibodies staining protocols. B, B', B'') Line profile across the center of the brain region for the images shown in Panel (A) (Dashed pink lines). C, C') Zoomed in visualization of the deepest region in the brain from the images shown in Panel (A) (Dashed pink boxes).

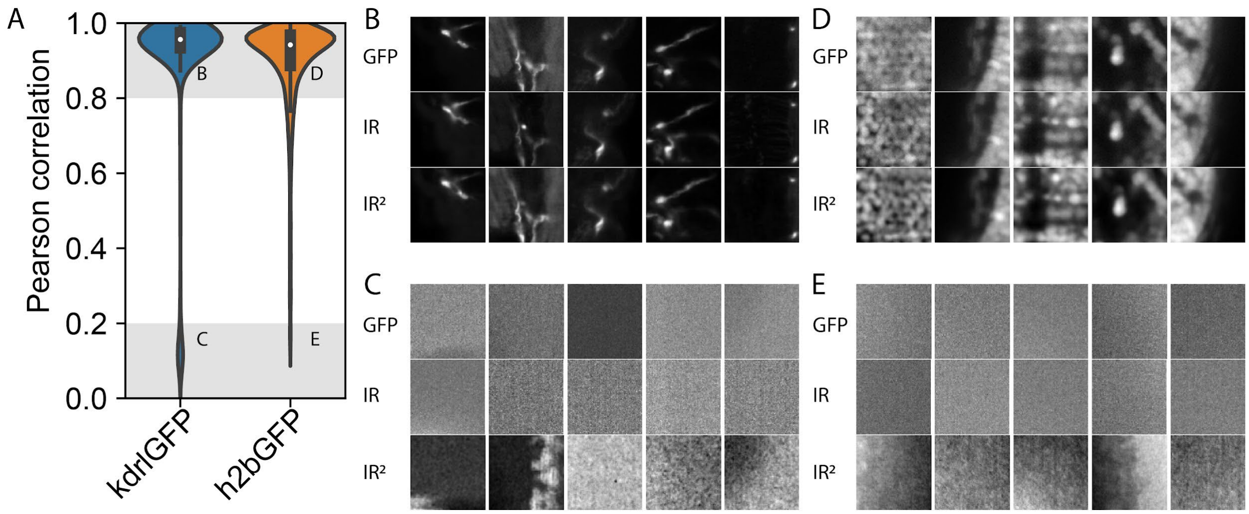


**Supplementary Figure 3.** A) Pearson correlation coefficient as shown in Figure 1 panel D. B, D) Patches examples with high correlation coefficients extracted from images of kdrl:GFP and h2b:GFP fish larvae, respectively. C, E) Patches examples with low correlation coefficients, extracted from images of kdrl:GFP and h2b:GFP fish larvae, respectively.

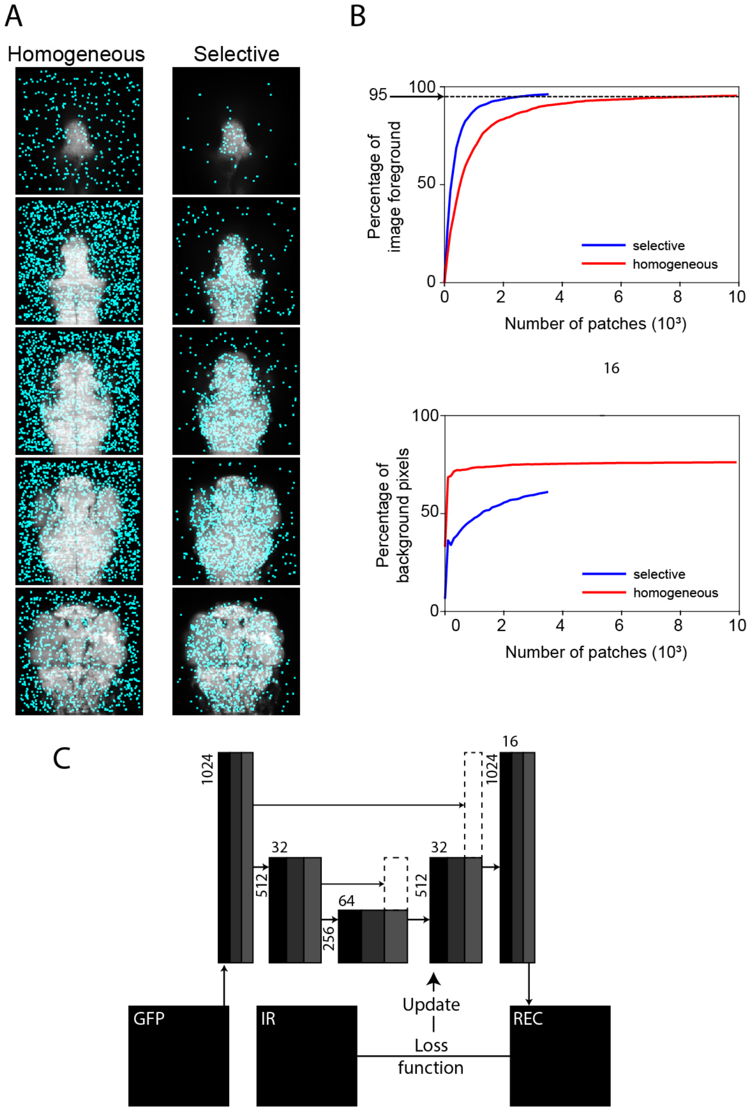


**Supplementary Figure 4.** A) Single Z planes of a fixed Tg(h2b:GFP) zebrafish larva (72 hpf). Transparent light blue square represents the patches extracted from the sample for network training. B) Fraction of sample coverage (top) and percentage of background pixels in the training set (bottom) with an increasing number of patches. Red and blue lines represent the fraction of pixels in high and low information content regions, respectively. C) A schematic of the U-Net deep learning network used for restoration. Horizontally and vertically aligned numbers denote the channel and spatial dimensions of the images at every layer of the U-Net architecture.

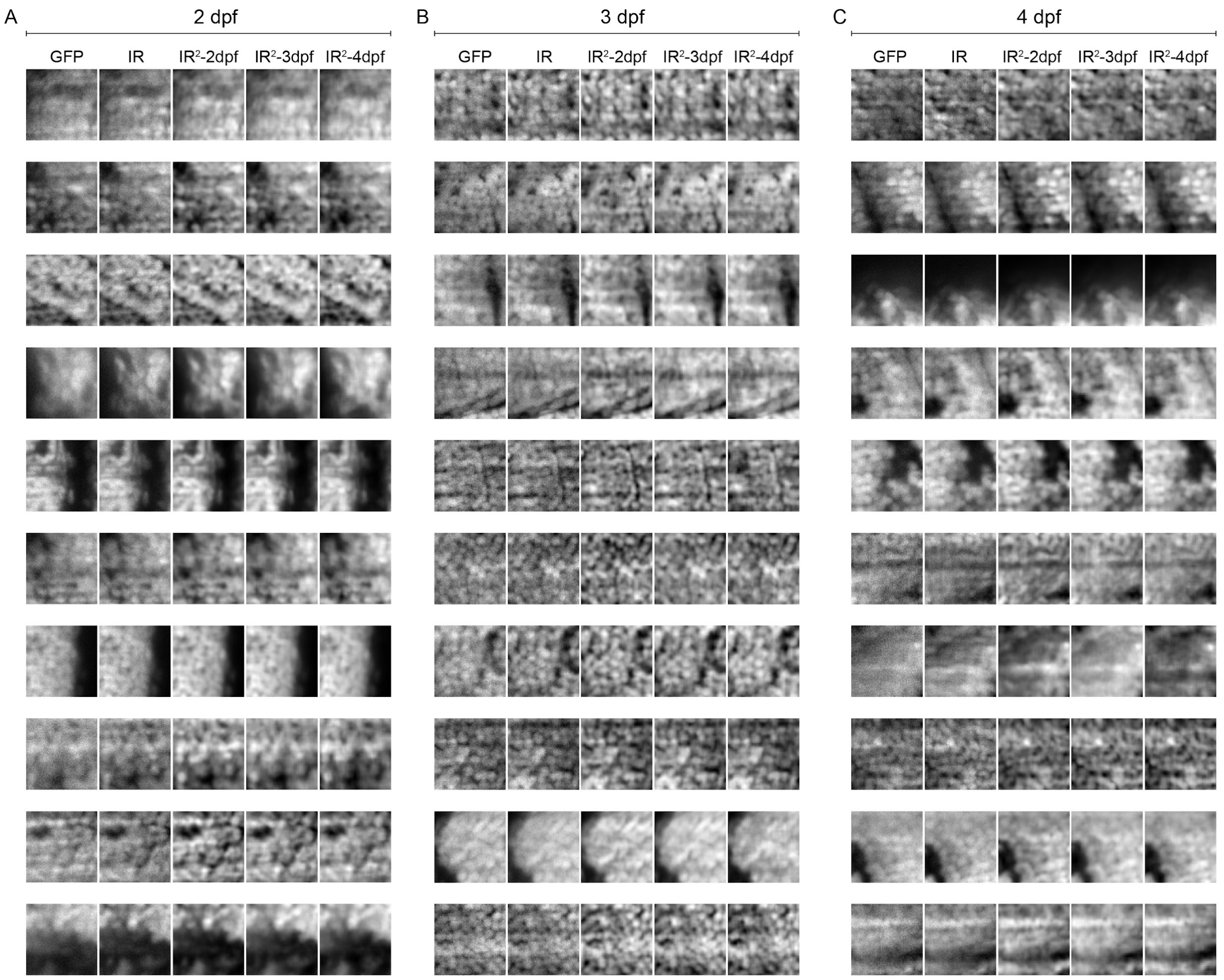


**Supplementary Figure 5.** Example patches from fish larvae images at 2 (A), 3 (B) and 4 © days post fertilization. First columns: input image (endogenous GFP), second column: infrared image, remaining columns: patches restored using models trained from fish at 2 (third column), 3 (fourth column) and 4 (fifth column) days post fertilization.

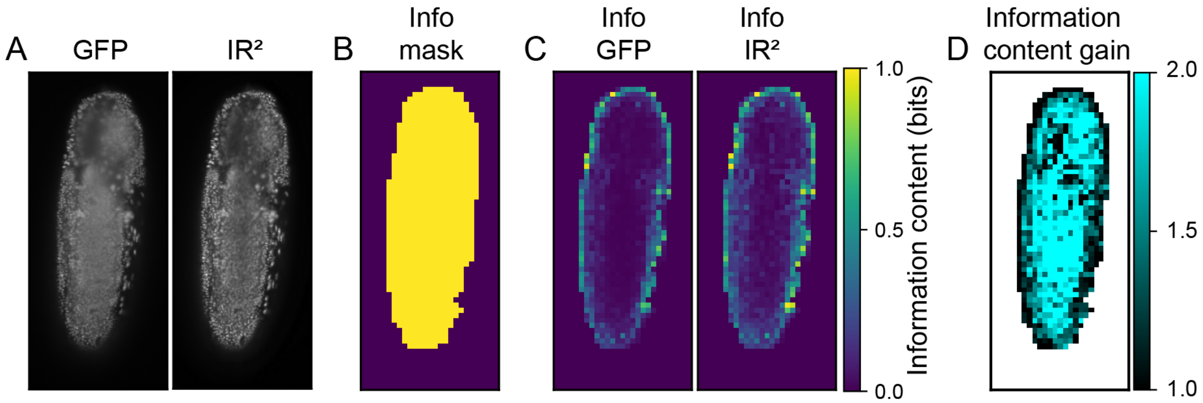


**Supplementary Figure 6.** Information content gain pipeline. A) Endogenous GFP and infrared individual Z-planes of a drosophila larvae used as input images. B) Binary mask obtained after automated thresholding. C) Absolute information content in the binary mask image, obtained using a sliding window. CD) Information content gain for the images shown in A), defined as the ratio between the information content of the images shown in C).
